# Generalized gametic relationships for flexible analyses of parent-of-origin effects

**DOI:** 10.1101/2020.04.13.039107

**Authors:** N. Reinsch, M. Mayer, I. Blunk

## Abstract

Genomic imprinting causes alleles to influence the phenotype in a parent-of-origin-specific manner. In attempts to determine the effects of imprinted loci, gametic relationship matrices have widely been used in pedigree-based parent-of-origin analyses of population data. One drawback of this is the size of these matrices because they represent each individual by two gametic effects. Significantly fewer equations are needed if a previously published reduced imprinting model is used that relates observations from progeny without its own offspring to the transmitting abilities of their parents. This can be accomplished using a numerator relationship matrix, with only a single row and column per parent and ancestors. However, the reduced model is not applicable when the parents have records. To better handle the curse of dimensionality, we propose a combination of average gametic effects (transmitting abilities) for individuals without their own records and single gametic effects for others. The generalized gametic relationship matrix is the covariance of this mixture of genetic effects that allows for a significant reduction in the number of equations in gametic models depending on the trait, depth of pedigree, and population structure. It can also render the reduced model much more flexible by including observations from parents. Rules for setting-up its inverse from a pedigree are derived and implemented on an open-source program. The application of the same principles to phased marker data leads to a genomic version of the generalized gametic relationships. The implementation of generalized gametic models to the ASReml package is illustrated through worked examples.

---

Shortly after its discovery, it was recognized that the gametic relationship matrix (Smith and Allaire, 1985; Schaeffer et al., 1989) can help isolate fractions of the genetic variance in quantitative traits caused by genomically imprinted loci. Alleles of the latter are expressed in a parent-of-origin-specific manner. In the early stages of pedigree-based imprinting analysis, animal models were augmented by an additional vector of paternal (alternatively, maternal) gametic effects, usually modeled as uncorrelated with any other effect. Its variance was assumed to be the product of a gametic relationship matrix and a variance component that can be explained by polymorphisms at loci with only paternal (maternal) gene activity. Pioneered by DeVries et al. (1994), these models were in use for more than a decade. However, they can account only for a single kind of classical imprinting, where either maternal or paternal alleles are fully silenced through, e.g., the methylation of DNA. A proposal (Hill and Keithly, 1988) to consider both kinds of imprinting simultaneously did not materialize in any pedigree-based analysis of empirical data. Further, there was uncertainty regarding ways to account for the effects of partially imprinted loci, where both alleles are expressed but at different strengths depending on their parental origins.

A model for parent-of-origin analysis was subsequently developed (Neugebauer et al., 2010a, b) that is comprehensive in the sense that it accounts for all kinds of imprinting, be it full or partial, maternal or paternal (Blunk et al., 2014). This so-called reduced imprinting model relates observations from non-parents (final progeny, e.g., animals used for meat) to transmitting abilities (half of the breeding values) of their parents. There are two correlated genetic effects per parent, a transmitting ability as sire and a transmitting ability as dam, which reflect an animal’s genetic effect on its offspring under a paternal or maternal imprinting pattern. In the presence of genomically imprinted loci, these two genetic effects are different. The variance of these differences has been called the imprinting variance because it summarizes contributions from all kinds of possible imprinted loci. A numerator relationship matrix is needed for parents only, as the final progeny with observations but without offspring do not appear in the underlying pedigree and the resulting relationship matrix. The null hypothesis of the absence of polymorphic imprinted loci with an effect on the trait under investigation (i.e., a zero imprinting variance) can be tested by a restricted maximum likelihood (REML) ratio test.

Alternatively to the above, a comprehensive *gametic model* can be used to estimate the same set of genetic covariances, including the imprinting variance (Tier and Meyer, 2012; Meyer and Tier, 2012). This requires four gametic effects to be estimated per individual, two as sire and two as dam, where the relationships include the final progeny with observations. As an advantage over the reduced model, the gametic model allows for records from parents. Moreover, it can be extended to account for maternal effects (see Appendix A5).

The use of measured genotypes in genomic best linear unbiased prediction models (gBLUP) that include imprinting effects has been outlined by Nishio and Satoh (2015). The first (GBLUP-I1) of the two variants of the proposed model contains an imprinting effect that is modeled as independent of the action of un-imprinted Mendelian locus, summarized as an additive genetic effect. The second model (GBLUP-I2) considers a paternal and a maternal gametic effect with zero mutual correlation. This clearly could be turned into a comprehensive model by abandoning the assumption of a zero correlation and replacing pedigree-derived gametic relationships by a genomic counterpart of equal size and structure. In cases where not all pedigreed individuals are genotyped, this enables a combined analysis of the genotyped and un-genotyped individuals in a single-step approach (Legarra et al., 2009; Aguilar et al., 2010; Christensen and Lund, 2010). The first model (GBLUP-I1), by contrast, cannot easily be extended to have such a pedigree-derived counterpart.

The downside of the gametic model is the large number of equations (Smith and Allaire, 1985) used to represent the random genetic effects, in particular when variance components are to be estimated. A pedigree with a size of approximately half a million is a technical barrier for REML estimation in animal models using currently available software packages (Shor et al., 2019). With a gametic parent-of-origin model, the same number of equations is reached with only a quarter of individuals. Therefore, the question arises if there is any option for models that retain the flexibility of the gametic model while allowing for a considerably smaller number of equations for random genetic effects, as close as possible to the reduced imprinting model.

As a solution, we propose a much smaller re-defined vector of genetic effects obtained by a proper linear transformation of the gametic effects. This is rendered applicable by introducing a version of a corresponding relationship matrix, called the generalized gametic relationship matrix, together with rules for its rapid inversion from the pedigree. As a result, the size of the gametic model can be reduced to a more manageable one while retaining all of its advantages. We also show how the same kind of transformation can be applied to measured genotypes to obtain conformable genomic and pedigree-derived versions of the new relationship matrix.

## THEORY

### Generalized gametic relationships

In gametic models, each individual *i* is represented by the additive genetic effects of its paternal gamete *g*_*i*,1_ and maternal gamete *g*_*i*,2_ (Schaefferet al., 1989), which usually are arranged in a pair-wise manner in a vector **g** of length 2*t*, which is twice the number *t* of individuals in the pedigree. The model equation for a phenotypic observation *y*_*i*_ of individual *i* then is

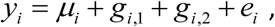

With 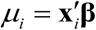 as a place-holder for any combination of explanatory variables in vector 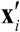 with fixed effects **β**, and the residual *e*_*i*_. Thus, the gametic model splits the additive genetic value (breeding value) *b*_*i*_ of individual *i* into paternally derived and maternally derived parts, *b*_*i*_ = *g*_*i*,1_ + *g*_*i*,2_.

The basic idea of reducing equations in gametic models by a considerable number is to replace the two gametic effects of a subset of *u* individuals by their pair-wise average:

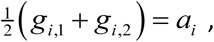

which is known as the transmitting-ability (half the breeding value) of individual *i*.

The vector **g** of gametic effects can be arranged such that the gametic effects of all *u* individuals precede the gametic effects of the *v* that are bound to retain their distinct gametic effects. The corresponding subdivision of **g** is

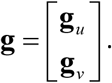

The sub-vectors **g**_*u*_ and **g**_*v*_ have respective lengths of 2*u* and 2*v*. The covariances of all gametic effects in **g** are the elements of the 2*t* × 2*t* gametic relationship matrix **G** (Schaeffer et al., 1989). It can be partitioned into sections that correspond to the relationships between the gametic effects in **g**_*u*_ and **g**_*v*_

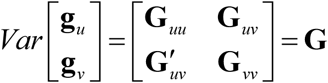

The required average gametic effects can be obtained by a linear transformation that is defined by a matrix **K**′, such that

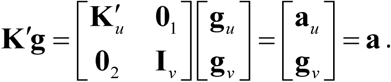

In effect, all gametic effects of individuals in **g**_*u*_ are replaced by their transmitting abilities in **a**_*u*_. The upper-left partition 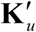 of the transformation matrix **K**′ has dimensions *u* × 2*u*, and is defined as the Kronecker product of a *u*×*u* identity matrix **I**_*u*_ and a row vector with two elements equal to 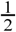:

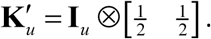

Further, **K**′ comprises a 2*v*× 2*v* identity matrix **I**_*v*_ and two null matrices, **0**_1_ and **0**_2_, with respective dimensions of *u* ×2*v* and 2*v*×2*u*.

The covariance matrix of the transformed vector of gametic effects **a** then becomes

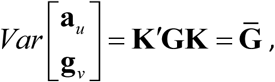

which in the following is called a generalized gametic relationship matrix. A natural choice is to retain the gametic effects of all individuals with their own phenotypes in vector **g**_*v*_ and let all their ancestors without records be represented by their transmitting abilities, constituting **a**_*u*_. The subdivisions of 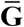 then are

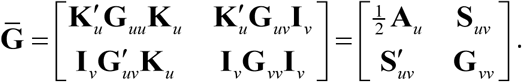

The upper-left part 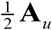 is equal to the co-ancestry matrix (half the numerator-relationship matrix) of all ancestors without own records, while **G**_*vv*_ reflects relationships between the gametic effects of all individuals with their own observations. Finally, **S**_*uv*_ contains the covariances between transmitting abilities and gametic effects. See the small example involving four individuals (IDs). There are three transmitting abilities for individuals 1, 2, and 3, with corresponding pair-wise elements of 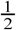 in the transformation matrix **K**′ and two gametic effects, for which the elements in **K**′ are one. The resulting generalized gametic relationship matrix 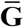 has dimensions 5×5.

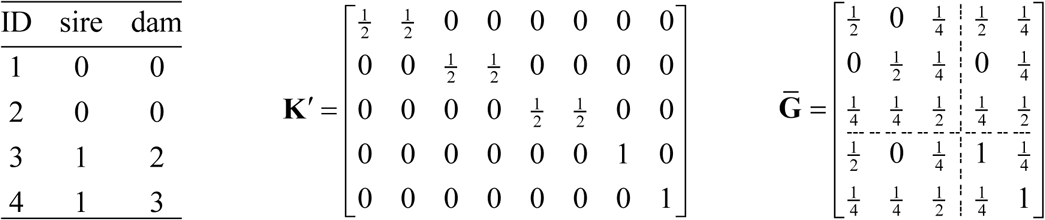

### Generalized gametic relationships in a gametic model

In light of the above, the model equation for an observation *y*_*i*_ can be retained as in the gametic model, and a mixed model that considers parent-of-origin effects (POEs) and uses the generalized relationship matrix becomes

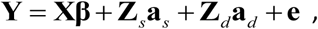

where **Y** is a vector of observations, **β** comprises the fixed effects, and **X** is the corresponding incidence matrix. The covariance of random effects is assumed to be

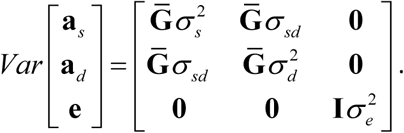

This generalized gametic model contains the gametic effect vectors **g**_*s*_ and **g**_*d*_ replaced by their transformed counterparts **a**_*s*_ and **a**_*d*_, respectively, and, consequently, uses the corresponding relationship matrix 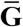 instead of the classical gametic relationships of **G**. Further, incidence matrices **Z**_*s*_ and **Z**_*d*_ link observations to the random gametic effects in **a**_*s*_ and **a**_*d*_, respectively, while no observation is linked to any of the transmitting abilities in the latter vectors. As a result, any incidence matrix **Z**^*a*^ = [**0**^*u*^ **Z**^*v*^] that links observations to gametic effects in the generalized vector of genetic effects 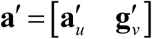 can be considered a converted incidence matrix **Z**^*g*^ = [**0**^2*u*^ **Z**^*v*^] from a classical gametic model that links the observations to the gametic effects in 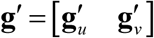:

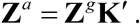

This transformation retains all columns in the partition **Z**^*v*^, i.e., one per gametic effect of individuals with records, while the number of null columns in **0**^*u*^ of **Z**^*a*^ collapses to half of that of **0**^2*u*^ in **Z**^*g*^. In the same manner, both incidence matrices **Z**_*s*_ and **Z**_*d*_ from the previous model equation are converted versions of their counterparts in the classical gametic imprinting model, which forms the basis for the proof of equivalence of the classical and the generalized gametic models involving 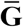 (see Appendix A1).

### Reduced gametic model

The reduced imprinting model as initially described by Neugebauer et al. (2010a, b) relates each observation from the final progeny *i* to the transmitting abilities as sire 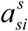 and as dam 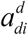 of the parents *si* (sire of i) and *di* (dam of i), respectively. For a single observation *y*_*i*_ we have the observation equation

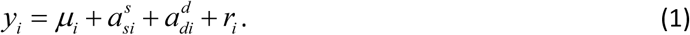

Here, the residual *r*_*i*_ is a sum of the Mendelian sampling effects of both parents (*m*_*si*_ and *m*_*di*_) and the measurement noise (*e*_*i*_). The latter is identical to the residual of the gametic model. Thus,

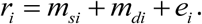

Its variance is a function of the inbreeding coefficients *F*_*si*_ and *F*_*di*_ of the parents of *i* :

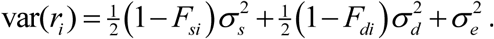

By rewriting the transmitting abilities of the parents as the averages of the respective gametic effects, i.e., 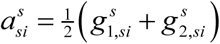 and 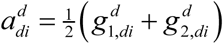, we get an observation equation in terms of gametic effects:

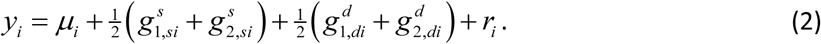

The covariance of the gametic effects then is

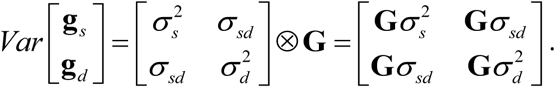

Here, the relationship matrix **G** of the gametic effects that define the involved transmitting abilities includes only the parents and their ancestors. The advantage of this gametic version of the reduced imprinting model over the previously published version that uses only transmitting abilities and their relationship matrix 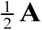 is that it enables us to easily integrate observations from parents by linking them to the respective gametic effects. Hence, for observations of any parent *i*, the observation equation becomes

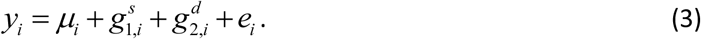

### Generalized reduced gametic model

The drawback of the reduced gametic model is that it has twice the number of equations compared to a version that uses 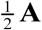. For all individuals without own records, it is however possible to reduce the number of equations for random genetic effects by representing the individuals through their transmitting abilities (average gametic effects) while retaining separate gametic effects for all parents with records, i.e., vectors of gametic effects **g**_*s*_ and **g**_*d*_ are replaced by appropriately transformed counterparts **a**_*s*_ and **a**_*d*_, respectively. Consequently the covariances of random genetic effects in a parsimonious generalized reduced gametic model that allows for parents with records is

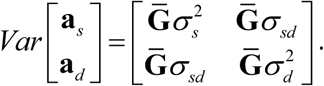

Further, we need a diagonal matrix **W** of weights equal to *w*_*i*_ = 1 for observations from parents, for which model Equation (3) applies and

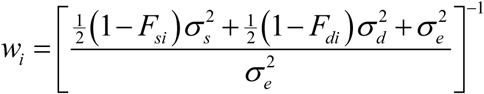

for the final progeny, where parents without their own records are represented by transmitting abilities or both parents have a record and are represented by gametic effects (the respective observation equations are (1) and (2)). The same weight applies to mixed kinds of representation that arise from cases where one parent of a final progeny has a record while the other does not. The corresponding observation equations for observations *y*_*i*_ of such final progeny are

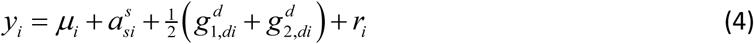

and

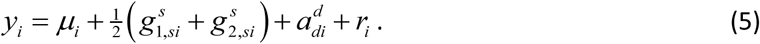

### A general model for parent-of-origin analyses

A general comprehensive model for parent-of-origin analyses banks on the generalized gametic relationship matrix. Special cases of the generalized gametic relationship matrix 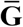 are the classical gametic relationship matrix 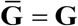 in the gametic model and 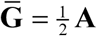 as in the reduced imprinting model. Correspondingly, the matrix **W** of weights can be an identity matrix that fits the classical gametic model, or a matrix all the weights of which are different from one as those in the reduced model for records of the final progeny. A general model can be specified for parent-of-origin analyses containing these two basic kinds of comprehensive imprinting models as well as models with any combination of gametic effects and transmitting abilities that can be obtained using our transformation matrix **K**′. In matrix notation, the general model is

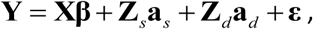

where **ε** is a vector of residuals. That is, *ε*_*i*_ = *e*_*i*_ for records from individuals represented by two gametic effects, or *ε*_*i*_ = *r*_*i*_ for observations from final progeny linked to the genetic effects of their parents. The respective weights are

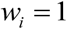

and

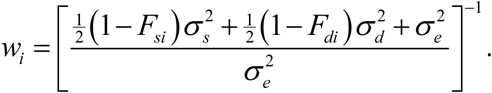

Random genetic effects and residuals are assumed to have covariance:

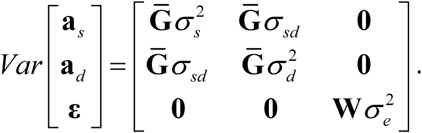

The resulting mixed model equations are

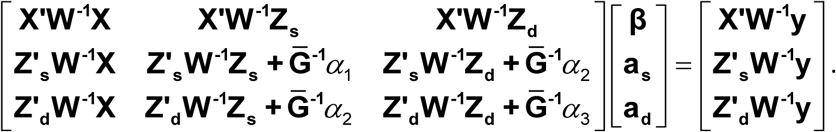

with

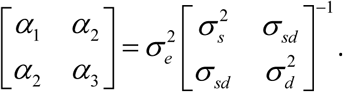

The general model comprehends any combination of observation Equations (1) to (5) to provide a large degree of flexibility in parent-of-origin analyses. Model variants may be chosen to minimize the number of equations for random genetic effect by using as many reduced observation equations as possible, which comes at the expense of the need for the recomputation of weights when estimating the components of variance. Alternatively, the repeated recomputation of weights may be avoided by representing all individuals with an observation using gametic effects. The underlying reason for this flexibility is that for the given data (observations, fixed effects, and pedigree), each possible general imprinting model has as an equivalent the same classical gametic model (that follows from Appendices A2 and A3). Consequently, any two general models that share the same equivalent classical model are also equivalent, and can replace each other, especially for the sake of estimating the components of variance.

### Direct inversion of the generalized gametic relationship matrix

Setting-up the inverse generalized gametic relationship matrix is key to any large-scale application. Rules for direct inversion can be derived by factoring the inverse 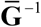 into inverses of a matrix **T**′ and a diagonal matrix **D** of inverse Mendelian sampling variances

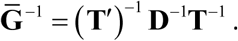

The above is known from the direct inversion of the numerator relationship matrix (Henderson, 1976; Quaas, 1976) and the classical gametic relationship matrix (Schaeffer et al., 1989). The matrix (**T**′)^−1^ is lower triangular, as shown in Figure 1. The underlying pedigree of this example (Supplement) comprises 12 individuals. Nine of them are represented by a single transmitting ability while the remaining three by two gametic effects. The kind of representation is indicated by the respective values of one and two in the last column of the pedigree file. Consequently, the dimensions of the inverse of the example are 15×15. Each of the 15 rows of (**T**′)^−1^ pertain to a single genetic effect, which itself may be derived from different kinds of genetic parental predecessor effects: An individual’s transmitting ability may be derived from two unknown parents (a-00) or a single unknown parent, where the known parent may be represented by a transmitting ability (a-0a, a-a0) or two gametic effects (a-0gg, a-gg0). Two known parents may show up as any combination of transmitting abilities or gametic effects (a-aa, a-agg, a-gga, a-gggg). Likewise, a gametic effect may be derived from an unknown parent (g-0), or a known parent enrolled by either a transmitting ability or two gametic effects (g-a, g-gg). These 12 cases need to be distinguished for directly inverting the generalized gametic relationship matrix. The example pedigree was constructed such that each case appeared at least once. For each effect related to a particular row of the lower-triangular matrix in Figure 1, the case is indicated in the last column. Note that the six cases a-0gg, a-gg0, a-agg, a-gga, a-gggg, and g-a are specific to the generalized gametic relationship matrix as they appear neither in the direct inversion of the numerator relationship matrix—involving only a-00, a-0a, a-a0, and a-aa—nor the classical gametic relationship matrix, for which only g-0 and g-gg need to be distinguished.

**Figure 1:**
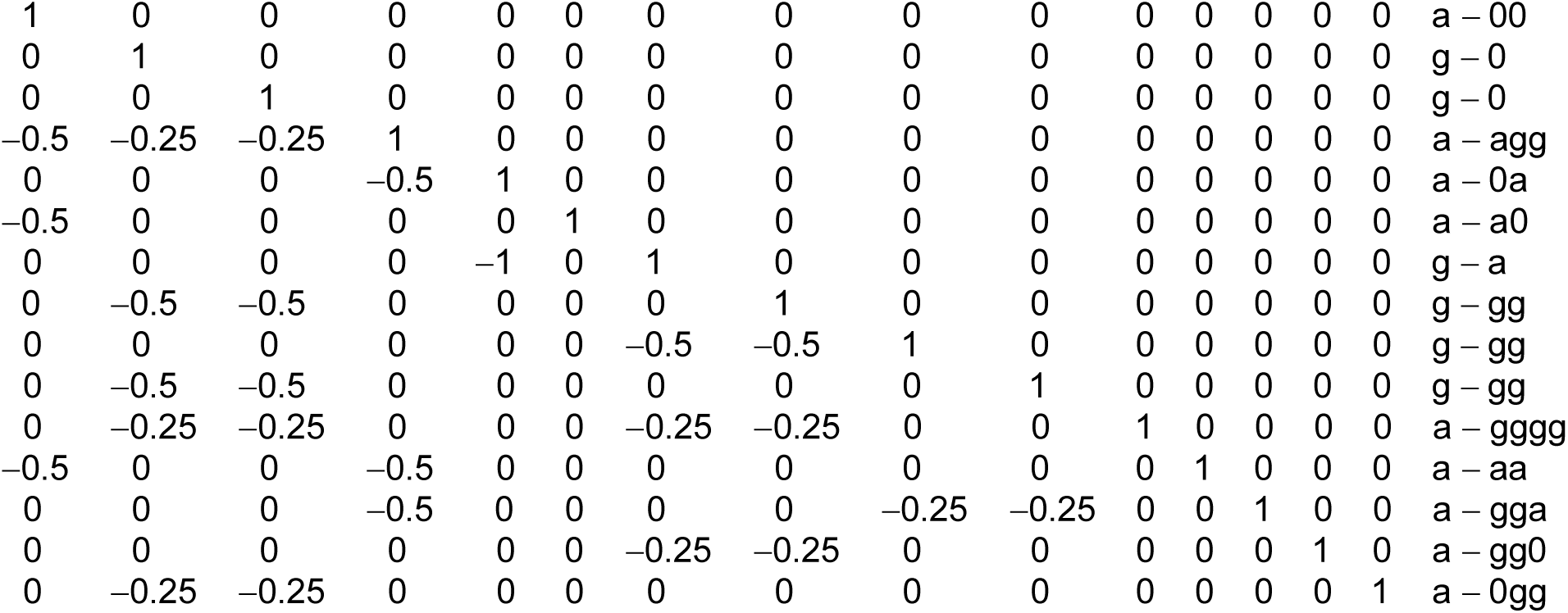
Example of a lower triangular matrix (**T**′)^−1^ from a decomposed inverse of a generalized gametic relationship matrix. Each row of the matrix pertains to a particular genetic effect. The last column indicates the respective combination of each kind of genetic effect (a: transmitting ability; g: gametic effect) with the genetic effects of the parents (a: transmitting ability; gg: pair of gametic effects; 0: unknown parent, and combinations thereof).

Mendelian sampling variances that define the diagonal elements of **D** are different for transmitting abilities and gametic effects. Further, they depend on the occurrence of unknown parents and the inbreeding coefficients of the known ones. In particular, this is *F*_known parent_ when an individual with a transmitting ability in the matrix has only one known parent, or *F*_sire_ and *F*_*dam*_ in case of full parentage information. For gametes, we need to account for the inbreeding coefficient *F*_parent_ of the known parent from which a gamete is derived. Accordingly, the 12 cases (a-00, a-0a, …, g-00) are grouped into five classes with distinct formulae for the inverse Mendelian sampling variance *δ* :

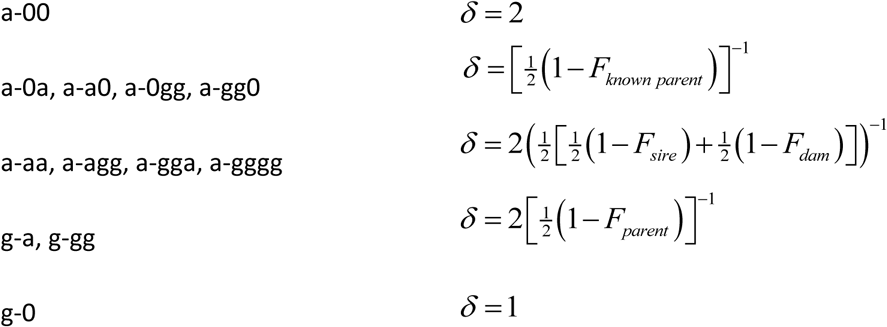

For any arbitrary order of genetic effects, the inverse generalized gametic relationship matrix can be constructed step by step from the pedigree. In each step, a matrix contribution **U**_*i*_ is added for genetic effect *i* to a matrix composed of an inverse 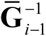 that already covers the preceding *i* −1 effects and zeroes:

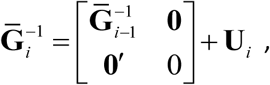

where **0** is a column vector of *i* − 1 zeroes and

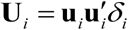

is the contribution made for each genetic effect *i*. The row-vector 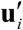 consists of all zeros, except for those elements with indices indicating the genetic effects of the respective parent(s). At minimum, the i-th element is always equal to unity as a non-zero element in this vector. All other non-zero elements are negative, with values of either 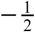 or 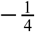. Thus, the number of non-zero entries varies from one to five, as can be derived from the rows of the example triangular matrix (**T**′)^−1^ in Figure 1. For all 12 possible cases, the non-zero coefficients in 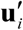 and their indices are summarized in Table 1. The non-zero elements of the resulting matrix 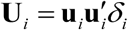 correspond to the (scaled) cross-products of the elements of the non-zero vector, and their coordinates in the matrix are the respective combinations of indices.

**Table 1:**
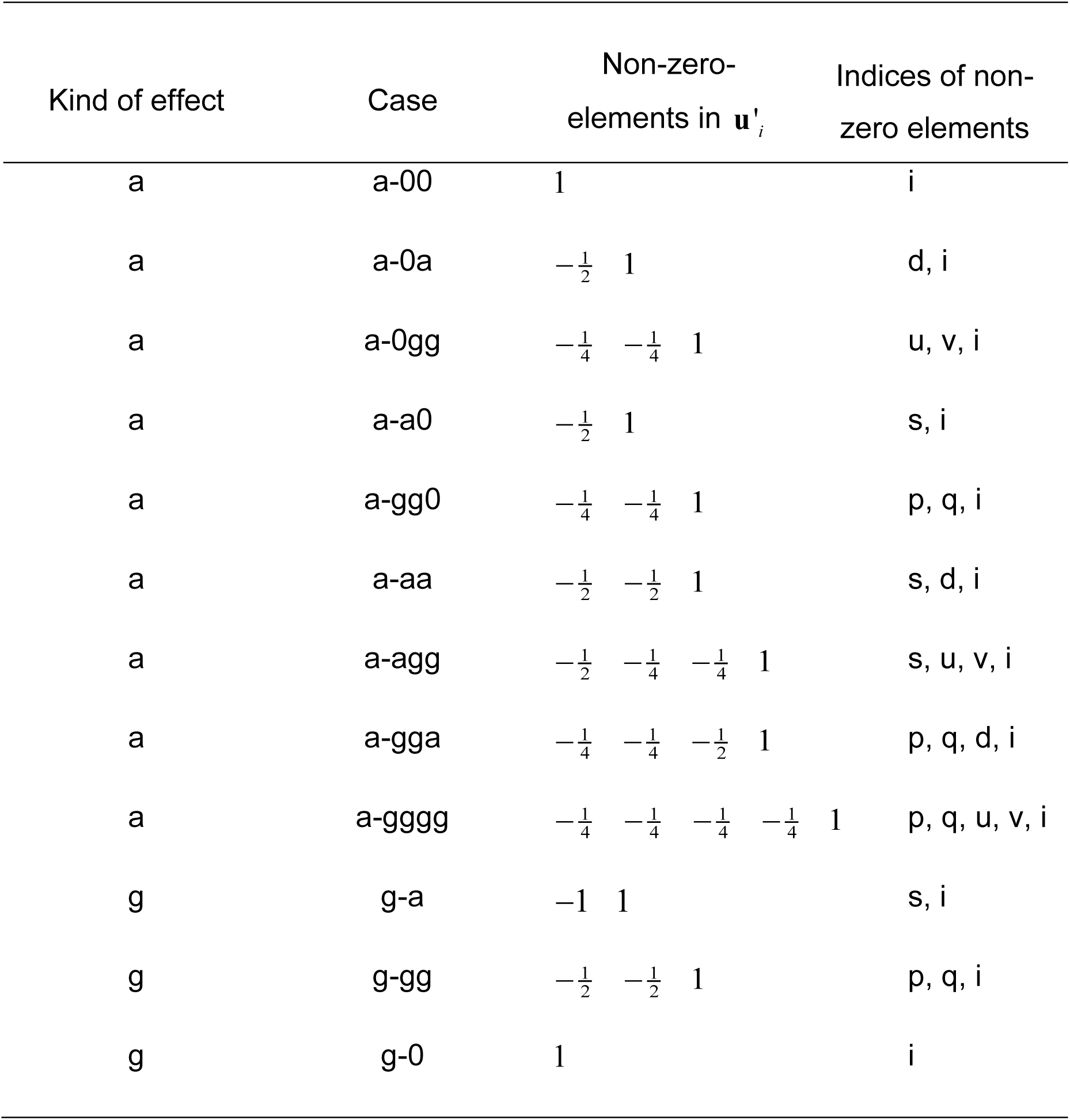
Size and indices of non-zero elements of vectors 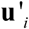 by kind of genetic effect (a: transmitting ability; g: gametic effect). The cases indicate unique combinations of kind of genetic effect and kind of indices. The latter consist of i: number of genetic effects; d: transmission abability of dam; s: transmission ability of sire; u: paternal gamete of dam; v: maternal gamete of dam; p: paternal gamete of sire; q: maternal gamete of sire. For gametic effects (cases g-a and g-gg), the respective effects of the known parent are indexed as for a sire.

| Kind of effect | Case | Non-zero-<br>elements in $\mathbf{u}'_i$ | Indices of non-zero elements |
| --- | --- | --- | --- |
| a | a-00 | 1 | i |
| a | a-0a | $-\frac{1}{2}$ 1 | d, i |
| a | a-0gg | $-\frac{1}{4}$ $-\frac{1}{4}$ 1 | u, v, i |
| a | a-a0 | $-\frac{1}{2}$ 1 | s, i |
| a | a-gg0 | $-\frac{1}{4}$ $-\frac{1}{4}$ 1 | p, q, i |
| a | a-aa | $-\frac{1}{2}$ $-\frac{1}{2}$ 1 | s, d, i |
| a | a-agg | $-\frac{1}{2}$ $-\frac{1}{4}$ $-\frac{1}{4}$ 1 | s, u, v, i |
| a | a-gga | $-\frac{1}{4}$ $-\frac{1}{4}$ $-\frac{1}{2}$ 1 | p, q, d, i |
| a | a-gggg | $-\frac{1}{4}$ $-\frac{1}{4}$ $-\frac{1}{4}$ $-\frac{1}{4}$ 1 | p, q, u, v, i |
| g | g-a | -1 1 | s, i |
| g | g-gg | $-\frac{1}{2}$ $-\frac{1}{2}$ 1 | p, q, i |
| g | g-0 | 1 | i |

### Transforming measured genotypes in a generalized genomic gametic relationship matrix

Parent-of-origin analyses may also use genomic relationships, or combined genomic and pedigree relationships. A specific feature of this is that ordinary marker genotypes (AA, AB, BB) are not sufficient for this purpose, and the parental origin of the marker alleles at each locus has to be inferred instead (Lawson et al., 2013, and references therein) and summarized as ordered genotypes AA, AB, BA, and BB, where the first allele is paternally derived. This is, however, not always possible for all members of a genealogy. In such a case, the principles above are beneficial for integrating ordered and unordered genomic information into a single genomic version of the generalized gametic relationship matrix.

Let us assume that all *t* individuals are genotyped with *p* markers and all genotypes are phased into 2*t* haplotypes. Information on the number (zero or one at each locus before centering) of minor alleles for all marker loci on each haplotype can be summarized in a column-wise mean-centered 2*t* × *p* matrix **C**. To this matrix, each individual *i* contributes two p-row-vectors 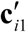 and 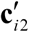, where the centered allele accounts for its first and second haplotype. Matrix **C** can then be split into two submatrices **C**_*v*_ and **C**_*u*_ :

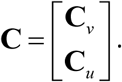

For imprinting analyses, at least all *u* individuals with records need to have their paternal and maternal haplotypes identified in **C**_*u*_. This can be achieved by adding at least one preceding generation without records but with genotypes. In case of only a single generation, all their 2*v* haplotypes in partition **C**_*v*_ would be left unordered. If the additional *v* genotyped individuals contain more than a single successive generation, only a part of their genotypes may qualify as ordered, with the exceptions coming from the founders.

From **C**, a genomic gametic relationship matrix can be derived:

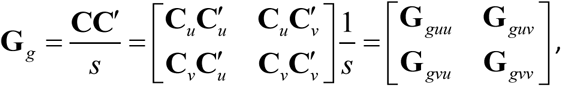

where *s* is a scaling factor, *s* = ∑ *p*_*j*_ (1 − *p*_*j*_), and *p*_*j*_ is the frequency of the allele at marker *j*.

In all cases where the parental origin of the two haplotypes can be traced back, the first haplotype of each individual is assumed to be paternal and the second maternal (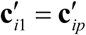 and 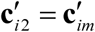); otherwise, the ordering of haplotypes is arbitrary. This is where the concept of generalization from above is used. A transformation matrix **K**′ can be defined such that for all individuals *i* with unordered genomic information, the two row vectors 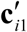 and 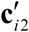 are replaced by their averages:

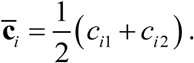

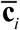 does not depend on the order or the parental origin of the haplotypes of an individual:

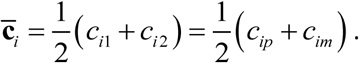

That is, 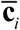 is also the vector of average paternal- and maternal-centered number of gene counts. Consequently, a generalized genomic gametic relationship matrix can be defined as

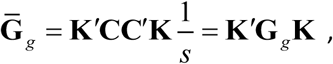

with **K**′ defined as before. The partition 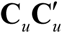 of **G** _*g*_ can be used to determine only the ordered genomic information of all individuals with records and, as such, is sufficient to estimate the components of genetic variance in a parent-of-origin analysis. All respective gametic effects of these individuals can also be estimated. The entire matrix **G** _*g*_ delivers gametic effects (as sire and dam) for all individuals, including those with no phenotypes. The generalized variant 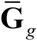 by design is also appropriate for parent-of-origin analyses, with no other requirements for **K**′ as for the pedigree-derived counterpart. Thus, the general model for parent-of-origin analyses is also applicable to genomic relationships, provided the marker haplotypes of individuals with observations can be ordered.

### Software and data availability

A detailed *guide to practical implementation* is available on the RADAR repository (https://www.radarservice.eu/radar/dataset/get/lGjshsdpCzWftGAQ?lang=en&token=DpsQlXcXJRuDkLmbwmzB – this is a temporary link for the purpose of review only and will later be replaced by a permanent DOI). It includes the source code of a program to directly set-up the inverse of the generalized gametic relationship matrix from a pedigree file, a detailed program description and example input and output files. There we also provide a collection of six worked toy examples demonstrating in very detail how various mixed models with generalized gametic relationships can be implemented using the ASReml package. Each example is also accompanied with R-code to check details and the correctness of the ASReml results.

## DISCUSSION

The outlined generalization introduces elements of the reduced imprinting model to the gametic model and vice versa, accompanied by gains in flexibility and substantial savings in terms of the number of equations used. The latter is important especially for estimating the components of variance (Shor et al., 2019). The matrix 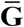 contains two limiting cases that set the boundaries for the ratio of equations that can be eliminated. The first is the classical gametic relationship matrix itself (dimensions 2*t* × 2*t*), when **K**′ is an identity matrix. The other limiting case is 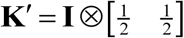, such that 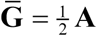 with dimensions *t* × *t*. Therefore, the reduction in the number of equations for genetic effects can take a range of 0%–50%, compared with a classical gametic model. However, the actual savings depend on the specifics of each dataset. As examples, two animal datasets were considered: the first was from an analysis of daily net gain in Brown Swiss fattening bulls (Blunk et al., 2018; Blunk et al., 2019), with a pedigree of 663,515 individuals (173,051 non-parents with records), whereas the second dealt with litter size in an experimental line of mice (2,137 females with an observation for first-parity litter size; necessery pedigree size for variance component estimation: 4544; total pedigree size 15222; unpublished data). In the Brown Swiss, the number of gametic effects for all animals was 1,327,030, compared with 836,566 with gametic effects for animals with observations only. The relative saving was 37% in terms of the number of equations and 32% in terms of the number of non-zero elements of the half-stored inverse. The respective numbers of equations in the mice example were 9088 versus 6681, with relative savings of 26% and 25%. In particular, small proportions of individuals that have records cause large reductions as all ancestors without a record are assigned only one equation. This applied to the mice example, as only females that had reproduced had records of litter size. If all available animals from the same number of generations were included (no “pruning” performed; 15222 animals), as one would prefer e.g. for the estimation of the genetic trend, there were 30,444 gametic effects versus 17,359 effects with the generalized relationships, with relative savings of 42% of equations and 56% of non-zero elements. The vast majority of this pedigree included males, females from older generations with no data, and non-reproducing females of younger generations.

Sex-specific traits such as litter size, number of eggs, or milk yield provide the opportunity to represent all males by their transmitting abilities. Thus, the resulting number of equations is considerably smaller in comparison with a trait recorded in both sexes. The family structure also has an effect: More equations are saved in the presence of typically small paternal groups of offspring, given that sires without own phenotypes are represented by their transmitting abilities. Further, it makes sense in imprinting analyses to add a high ratio of ancestors without phenotypes to better reflect inbreeding, and the relationships between genetic effects as sire and dam. Including their transmitting abilities rather than gametic effects in the model therefore also leads to a large number of saved equations.

A certain fraction of individuals with records might have either not reproduced at all or not yet reproduced at the time of data recall (i.e. they appear as final progeny), which provides the opportunity for representing them by reduced observational equations rather than having their own gametic effects in the model. In the Brown Swiss example, where all observations were from final progeny, this leads to a fully reduced model with relationships of 490,464 ancestors, a reduction of 63 %. In the small mouse example more equations for 634 final progeny can be saved (3910 animals and 5413 equations left), forcing the relative savings up to 40% of equations.

In certain cases, one could, however, abstain from reduced observational equations, which has the advantage that no weights are required that depend on as-yet undetermined components of variance. That has not proven to be a particular problem in the REML estimation of the components of variance (Neugebauer et al., 2010a, b; Blunk et al., 2017a,b), but may be beneficial to avoid in Bayesian approaches that employ Markov chain Monte Carlo methods, where the values of the components of variance change from iteration to iteration. By capitalizing on the flexibility of the generalized approach, weights become obsolete by representing all individuals with records—be they final progeny or not—by two gametic effects, which helps offset the computational burden resulting from repeated reweighting. At the same time, individuals without observations can be integrated by single equations.

For reasons of principle, a maternal genetic component of variance provides a special challenge as it is difficult to separate from the imprinting variance. Okamoto et al. (2019) showed that when estimated with a model variant that uses information only on the sire and maternal grandsire (Blunk et al., 2017; Okamoto et al., 2019), the imprinting variance may also be interpreted as maternal genetic. Similarly, for the reduced imprinting model, it can be shown that the imprinting variance and maternal genetic variance cannot be disentangled when both are present, and instead only a composite component of variance can be inferred (Appendix A4). A way out of this is to avoid reduced model equations and, instead, to represent individuals with records explicitly by their gametic effects in a model that includes maternal genetic effcts. Then, gametic variances as sire and dam can, at least in principle, be separated from the maternal genetic variance (Appendix A5). In practice, however, this may be hampered by limitations in the amount and structure of the data, as has been reported for Mendelian models (Heydarpour et al, 2008). Like maternal effects models, other kinds of imprinting models may also comprise more than a single genetic effect as sire and dam per individual—e.g., random regression models or multitrait models. As they all suffer from a large number of gametic equations, they benefit even more from generalized relationships.

In applications where all *v* individuals with records plus at least one preceding generation have measured genotypes and variance componets are to be estimated, it is sufficient to include only the subset of these *v* individuals with their genomic covariance **G** _*gvv*_. If there is interest in the genetic effects of the *u* founders as sire and dam, either **G** _*g*_ or 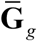 is the choice. An example is an F_2_ line-cross experiment with phenotypes recorded only in the F_2_ generation, and the genotypes of F_1_ and P_0_ generations needed only for phasing and determining line origins of the markers.

Often in animal breeding, large pedigrees are combined with smaller cohorts of genotyped individuals. Then, certain individuals are the first in their genealogy to be genotyped while the pedigree can be traced further back. In contrast to their own descendants, haplotypes of such a candidate cannot be ordered, which renders uncertain whether the first of two unordered marker haplotypes matches the paternal gametic effect in a pedigree-derived gametic relationship matrix or the maternal one. Consequently, a combined relationship matrix that is suitable for parent-of-origin analyses cannot be constructed. This problem can be solved by collapsing gametic effects into transmitting abilities both in the genomic relationships and the pedigree-derived ones. Then, generalized pedigree relationships for all animals can be combined with their matching generalized genomic counterparts 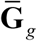 for the genotyped cohort in a way that allows for the easy integration of unordered genomic information. To this end, the available theory (Legarra *et al.* 2009; Christensen and Lund 2010; Aguilar *et al.* 2010) can be used to combine pedigree-derived relationships (here, 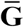) and genomic relationships 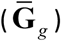 into a joint matrix, at least in the many cases where candidates with unordered genotypes have no record, such as dairy bulls.

In conclusion the generalized gametic relationship matrix provides the necessary flexibility to adapt <u>imprinting</u> analyses to specific computational and analytical needs in a large variety of situations through tailored versions of the general imprinting model. The most important aspects are the effective estimation of the imprinting variance in REML and Bayesian approaches in case the parents have records and the inclusion of maternal genetic effects and genomic relationships that integrate ordered and unordered genomic information. All things considered, these new possibilities are expected to stimulate systematic research on the importance of parent-of-origin effects for the genetic variation of quantitative traits in farm animals and other species.

## Acknowledgements

Inga Blunk was was funded by the Deutsche Forschungsgemeinschaft (DFG, German Research Foundation) project 418890112 “Making unused data resources available for imprinting analyses by using new methods to uncover parent-of-origin effects in human and livestock”.

## APPENDIX

### Appendix A1: Equivalence of the classical gametic model and the generalized gametic model in which all individuals with records have two gametic effects

Both models have the same expectation *E* (**y**) = **Xβ** of the vector of observations **y**.

The variance of observations in the classical gametic model is 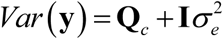,
where

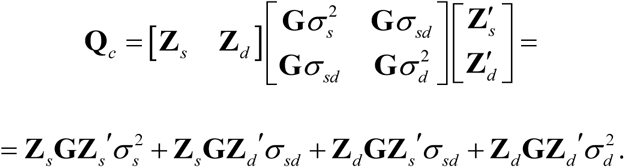

The first term can be rewritten as

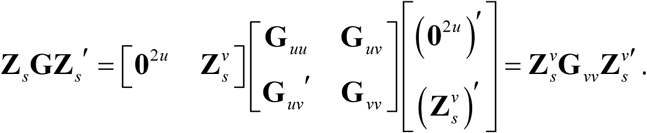

Likewise,

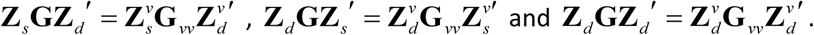

Finally,

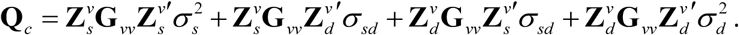

In the generalized case, the variance of observations is

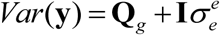

with

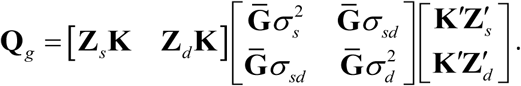

We make use of 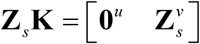 and 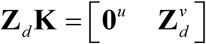, and rewrite

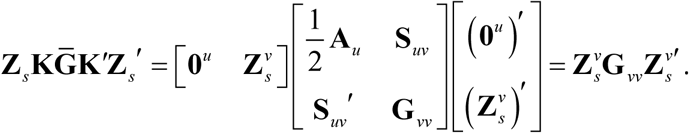

In the same manner,

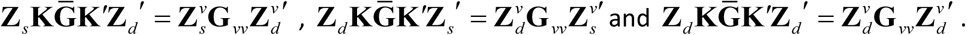

From this, we get

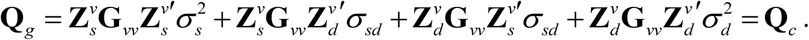

From **Q**_*g*_ = **Q**_*c*_, it follows that *Var* (**y**) is the same in both models and that they are equivalent.

### Appendix A2: Equivalence of classical and generalized gametic relationships in reduced models

We consider a reduced model with classical gametic relationships. With classical gametic relationships all parents of final progeny and their ancestors have two gametic effects in the model with covariance **G**. In the generalized case the gametic effects of *u* of them are collapsed into transmitting abilities, while the remaining *v* individuals retain their gametic effects. For the sake of generality the latter group, among an arbitrary choice of others, includes all parents who may have records. Parents with records need to have gametic effects, while all other individuals may be modelled by gametic effects or by transmitting abilities. As final progeny have no genetic effects of their own in the reduced model the variance of residuals 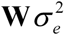 is not affected by relationships. With classical gametic relationships the variance of observations is

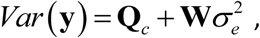

where

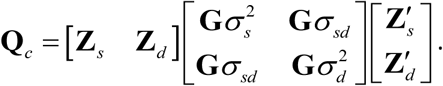

The incidence matrix **Z**_*s*_ for genetic effects can be partitioned as 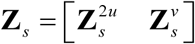. In the first partition are two adjacent columns per individual, i.e.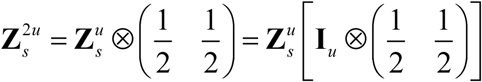, where 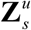 is the corresponding partition from the same kind of incidence matrix in the model with generalized gametic relationships and **I**_*u*_ is an *u* × *u* idendity matrix. Note that a multiplication with **K** cannot apllied here for the conversion of the matrix 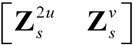 into 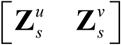, because 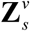 may both have entries of single ones for records of parents and of pairs of one half for records from final progeny. All of that also applies in an analogous manner to 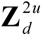 and 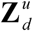.

The first component of **Q**_*c*_ is

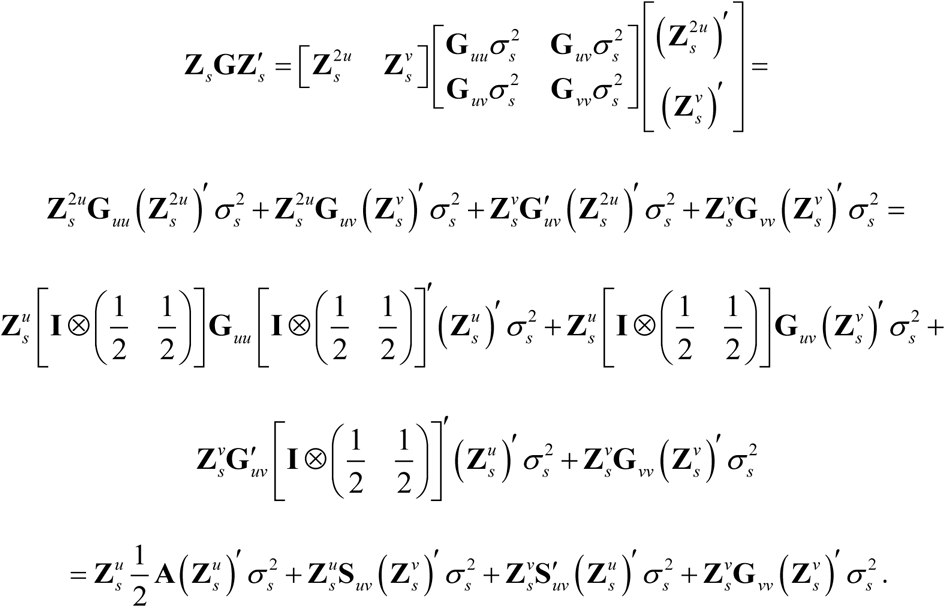

Similarly,

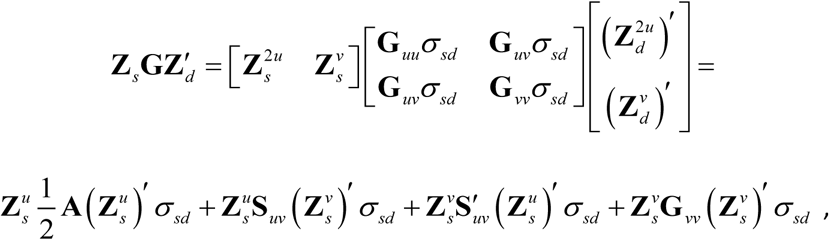

and

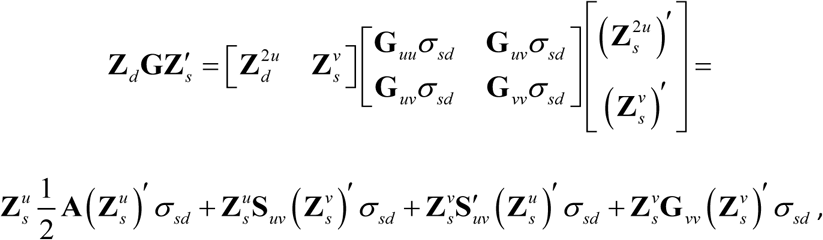

finally

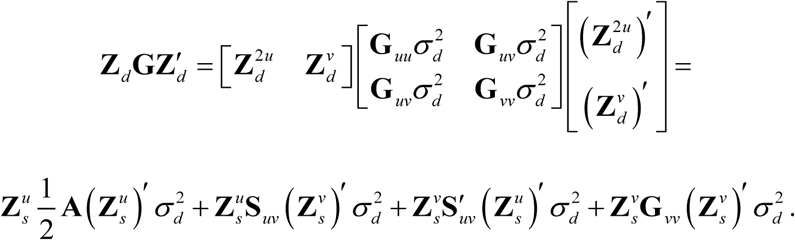

From this, **Q**_*c*_ can be summarized as

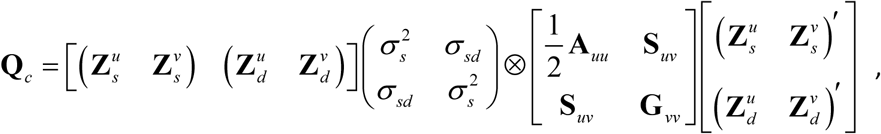

which is equal to the equivalent quantity **Q**_*r*_ using generalized gametic relationships

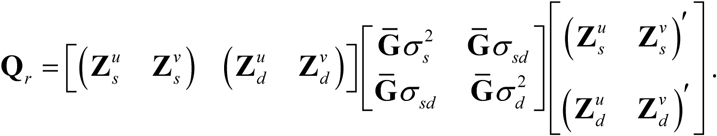

Thus

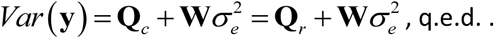

### Appendix A3: Equivalence between the model with gametic effects for all individuals and and the reduced model with gametic effects for parents

We consider a classical gametic model that includes a number *f* of final progeny. The vector **g**′ is partitioned into two components; in 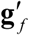 are the 2 *f* gametic effects of the final *f* progeny, and other gametic effects are in 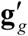. The covariance of **g**′ then is

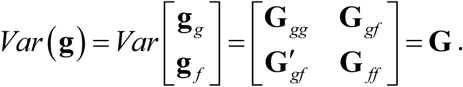

The incidence matrices for gametic effects are

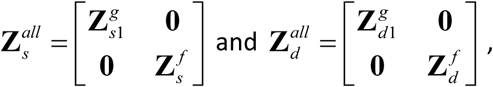

where 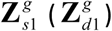 relates the observations to the gametic effects as sire (as dam) of individuals that are not in the set of the *f* final progeny. Accordingly 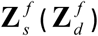 relates observations of the *f* final progeny to their respective gametic effects as sire (as dam).

By contrast, in a reduced model, all observations of the *f* final progeny are to be related to the gametic effects as sire (as dam) of their parents. The respective incidence matrices are

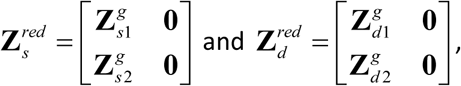

where 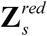 and 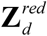 have only zero entries in columns for gametic effects of final progeny.

The relationships between the incidence matrices of the two types of models are

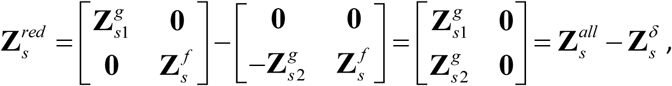

The matrix 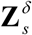 is the difference between 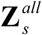 and 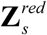:

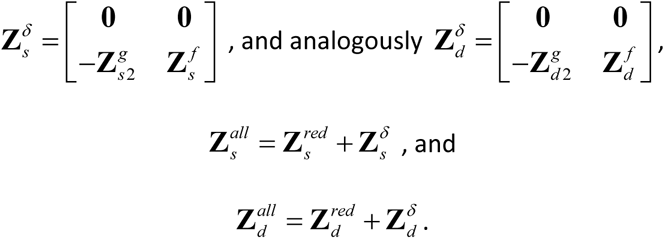

For the proof of equivalence of the two models, we express the variance of observations *Var* (**y**) in terms of model-specific incidence matrices and show their equality by making use of the last two identities.

For any reduced observation equation the variances of the relevant Mendelian sampling effects are part of the residual. For each paternal gamete as sire of a final progeny, the Mendelian sampling effect is the difference between the effect of the paternal gamete and the transmitting ability of the individual’s sire as sire. The respective vector is

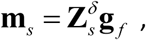

and the maternal counterpart as dam is

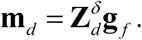

The common covariance matrix is

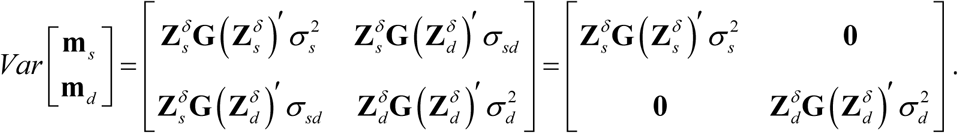

The covariance between **m**_*s*_ and **m**_*d*_ is zero as all rows of 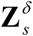 have their non-zero entries at places other than the rows of 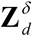, causing 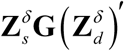 to be a matrix of zeroes.

In detail the product is

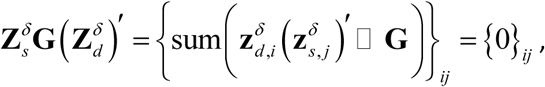

where □ denotes element-wise multiplication, sum () is the sum of all matrix elements in (), and 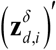 and 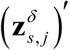 are the *i* th and *j* th rows of the two involved incidence matrices.

The total *Var* (**y**) in the reduced model is

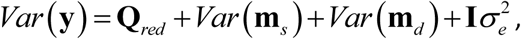

with

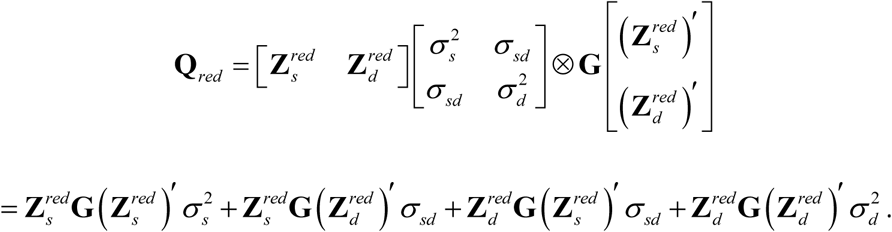

In the classical gametic model the variance of observations *Var* (**y**) is

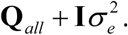

The first component is

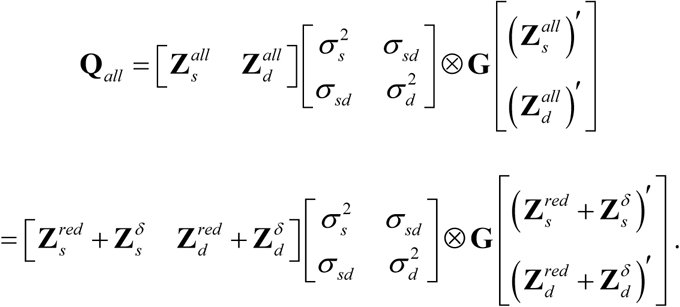

This results in a sum of 16 terms, of which the first four are

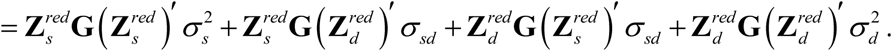

This is equal to **Q**_*red*_ in the reduced model. Further, we have two more terms

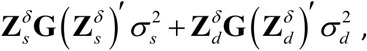

equivalent to *Var* (**m**_*s*_) +*Var* (**m**_*d*_). The remaining 10 terms in

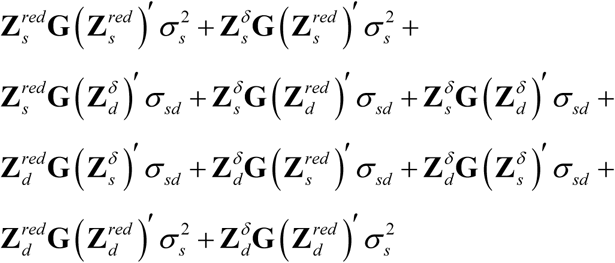

all are zero matrices. Thus, **Q**_*all*_ = **Q**_*red*_ +*Var* (**m**_*s*_) +*Var* (**m**_*d*_) and, therefore, the variance *Var* (**y**) in the classical gametic model and the reduced model with gametic relationships are identical. As both models also have identical expectations of **y**, they are equivalent; **q.e.d**..

### Appendix A4: Maternal genetic variance in a reduced model

We consider a reduced model equation for a single observation:

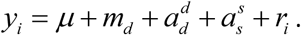

This equation comprises the maternal breeding value *m*_*d*_ of the dam *d* of individual *i*, together with the transmitting ability as dam 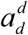 of the dam *d* of *i*, the transmitting ability as sire 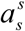 of the sire *s* of *i*, and the residual *r*_*i*_.

Then, the covariance of the respective vectors of random genetic effects is

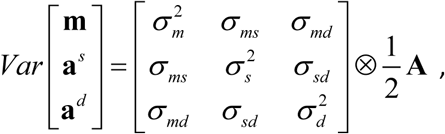

where 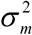 is the maternal gametic variance. As we use 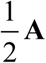 as relationship matrix (assuming all records are from final progeny) the incidence matrix for maternal breeding values needs to have non-zero entries of two to match this set of covariances.

The variance of observations has the non-residual component

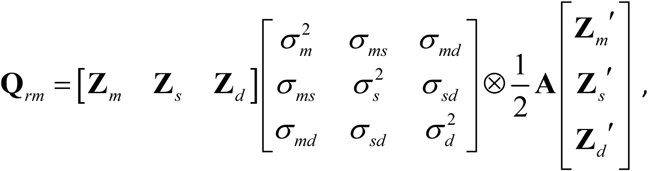

involving the incidence matrices **Z**_*m*_, **Z**_*s*_, and **Z**_*d*_ that link observations to maternal genetic effects, transmitting abilities as sire, and transmitting abilities as dam, respectively. **Q**_*rm*_ is a sum of nine matrices; of them, the following matrix equalities can be found by dropping the respective components of variance:

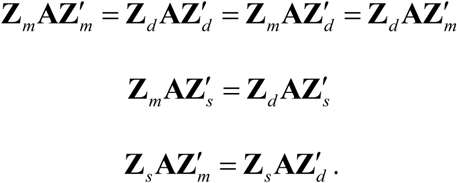

The underlying fact is that the incidence matrices **Z**_*m*_ and **Z**_*d*_ link all observations to genetic effects of the same animals, i.e., of the dam of each final progeny. Thus, the incidence matrices **Z**_*m*_ = **Z**_*d*_ are equal, and constitute equalities from above. Consequently, **Q**_*rm*_ can be rewritten as

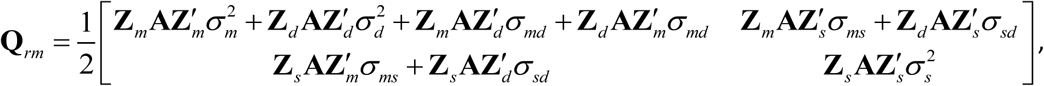

which, in terms of the incidence matrices of the reduced model without maternal genetic effects, is

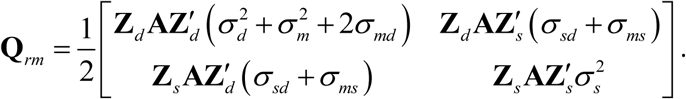

The variance in the transmitting ability as dam and the covariance with the transmitting ability as sire are therefore contaminated by components of the maternal genetic (co-)variances. This shows that in the presence of maternal genetic effects, 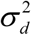 and 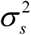 cannot be inferred from the reduced model. Moreover, we cannot correctly calculate the weights of the observations as this would require that we know these two components of variance.

Interestingly, we can assume the absence of genomic imprinting and make use of 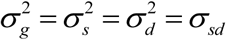, from which the residual variance of observation *i* becomes

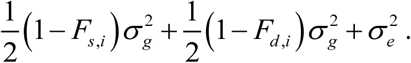

Consequently, the imprinting variance becomes

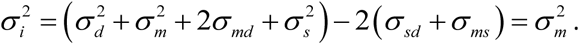

### Appendix A5: Maternal variance in a classical gametic model

The model equation for a single observation *y*_*i*_ in a gametic model with maternal effects is

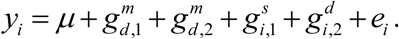

In this, we have the maternal effect (superscript *m*) of the paternal (1) gamete 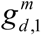 and the maternal (2) gamete 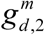 of the dam *d* of individual. 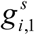 is the effect of the paternal gamete of individual *i* as sire, 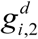 is the effect of the maternal allele of individual *i* as dam, and *e*_*i*_ is the residual. The covariance of random gametic effects is

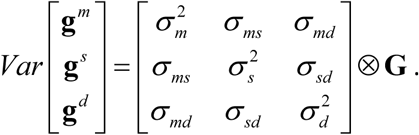

The vector of observations **y** has covariance 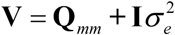, with

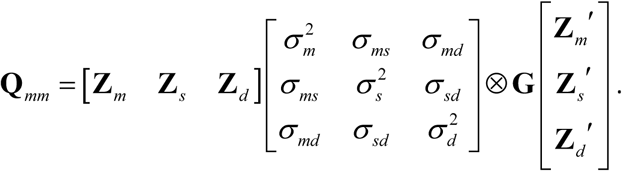

All other components of covariance are defined as before in appendix A4.

**Q**_*mm*_ has three components 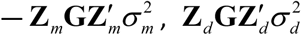, and —related to components of variance, and another three— 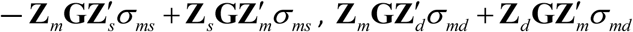, and—that are connected to the covariances. In contrast to the reduced model, the incidence matrices **Z**_*m*_ and **Z**_*d*_ relate the records to different gametic effects and, therefore, are not equal. As a result, all six addends of **Q**_*mm*_ are linearly independent and all components of variance can be separated.

